# Antigen Stimulation Reactivates HIV-1 Proviruses Despite Integration in Repressive Chromatin

**DOI:** 10.64898/2026.06.11.731680

**Authors:** Angelica Camilo-Contreras, Filippo Dragoni, Hao Zhang, Junlin Zhuo, Daniel Smith, Arturo Casadevall, Jun Lai, Pablo Tebas, Janet D. Siliciano, Robert F. Siliciano, Stuart Ray, Luis J. Montaner, Joel N. Blankson, Francesco R. Simonetti

**Affiliations:** Johns Hopkins University, Department of Medicine, Division of Infectious Diseases, Baltimore, MD, USA; Johns Hopkins Bloomberg School of Public Health, Department of Molecular Microbiology and Immunology, Baltimore, MD, USA; HIV Cure and Viral Diseases Center, The Wistar Institute, Philadelphia, PA, USA; Penn Clinical Trials Unit, University of Pennsylvania, PA, USA

**Keywords:** HIV-1, Quantitative Viral Outgrowth Assay, Viral Latency, Viral reactivation, Antigen Stimulation, Repressive Chromatin

## Abstract

Intact HIV-1 proviruses become progressively enriched in transcriptionally repressive genomic regions during long-term antiretroviral therapy (ART) and in elite controllers, raising questions about their capacity for reactivation in vivo. We used an antigen-restricted quantitative viral outgrowth assay (ag qVOA) to test whether cognate antigen stimulation can reverse latency of proviruses integrated within repressive chromatin. Using cells from two people with HIV (PWH) on ART, one on long-term treatment and one an elite controller, we show that antigen-specific stimulation induces viral outgrowth from intact proviruses integrated into a pericentromeric transition region and a zinc finger gene, respectively. These findings demonstrate that antigen recognition can overcome epigenetic constraints to reactivate proviruses with low inducibility and suggest that proviruses in so-called “deeper latency” may contribute to residual viremia and viral rebound following treatment interruption.

## INTRODUCTION

The Human Immunodeficiency Virus type 1 (HIV-1) infects CD4⁺ T cells by integrating its genome as a provirus into the target cell’s DNA. Antiretroviral therapy (ART) effectively blocks viral replication, leading to a rapid decline in the frequency of infected cells^1–3^. Despite this, a small reservoir of cells harboring replication-competent proviruses persists life-long due to clonal expansion^4,5,6,7–10^ and drives rapid viral rebound after treatment interruption^11,12^. Defining the mechanisms that govern viral rebound is central to HIV-1 cure strategies, yet the physiological triggers of latency reversal in vivo remain incompletely defined^13^.

Antigen-driven activation of CD4^+^ T cells maintains the reservoir by expanding infected clones^14–16^. Whether antigen recognition can also directly reverse latency in vivo remains less clear. Prior ex vivo studies from our group demonstrate that cognate antigen engagement can induce HIV-1 transcription^17,18^. In vivo, physiological signals can drive low-level residual viremia even under suppressive ART^19,20^, and responses to self-antigens have been associated with non-suppressible viremia^21^ (NSV).

The ability of a provirus to be reactivated depends in part on its integration site, which is itself determined by several layers of virus-host interaction^22^. Although the HIV-1 intasome favors bodies of actively expressed genes^23–25^, intact proviruses become progressively enriched within repressive chromatin regions, including zinc finger genes, lamina-associated domains, and pericentromeric zones. This pattern is observed in elite controllers^26,27^ (ECs, individuals who suppress HIV-1 replication without ART), individuals on long-term ART (LT-ART), and post-treatment controllers^28,29^. These findings suggest immune-mediated selection favors proviruses in repressive chromatin and that these proviruses reside in “deeper latency” due to their integration sites, allowing them to evade immune clearance^17,30^. However, integration within these regions does not uniformly confer transcriptional silencing, as reactivation potential depends on local chromatin features rather than integration site category alone^31,32,33^, as exemplified by a highly reactivatable clone in a pericentromeric region on chromosome 22^34^. Additionally, the inducible reservoir as measured by the quantitative viral outgrowth assays (qVOA) does not decline despite decades of immune selection on ART and even proviruses in zinc finger genes can be reactivated^4^, leaving the clinical relevance of “deeper latency” unresolved.

Based on evidence that antigenic stimulation can induce HIV-1 gene expression, we hypothesized that exposure to commonly-encountered antigens could trigger viral outgrowth, even from proviruses in repressive chromatin. We developed an antigen-restricted qVOA (ag qVOA) and determined that stimulation with cognate antigens can reverse latency ex vivo. Using cells from an individual on LT-ART and an EC, we show that stimulation with *Candida albicans* (CA) and HIV-1 Gag antigens induce robust viral outgrowth from proviruses integrated in transcriptionally repressive regions.

## RESULTS

### Antigen-specific stimulation induces minimal CD4⁺ T cell activation compared to polyclonal

### stimulation

The qVOA remains the gold standard to measure the minimal frequency of cells carrying inducible, infectious proviruses^35^. Despite recent advancements in quantifying intact proviruses via PCR^36,37^, the qVOA remains the most specific approach to study proviruses that rekindle viral replication if ART is removed. The qVOA relies on polyclonal T cell activation via PHA and allogenic stimulation with irradiated feeders, which maximally activates T cells but does not permit focused stimulation of CD4^+^ T cells reactive to antigens of interest. Importantly, it may not reflect the nuances downstream of the highly regulated TCR-peptide-MHCII driven activation.

To address these limitations, we developed the ag qVOA, which couples exposure to defined antigens with quantitative measurement of replication-competent virus, enabling assessment of latency reversal driven by antigen recognition (Figure 1A-C). We stimulated CD8^+^ T cell-depleted peripheral blood mononuclear cells (PBMCs) rather than isolated resting CD4^+^ T cells to permit ex vivo antigen presentation, replaced cytokine-enriched “Super T cell media” with a minimally activating medium containing costimulatory antibodies (CD28/CD49d) and IL-2 (10 U/mL), and omitted irradiated feeder cells from all conditions. As in the standard (std.) qVOA, cells treated with the mitogen phytohemagglutinin (PHA) were stimulated for 16 hours to avoid cytotoxicity. In contrast, cells treated with the antigen of interest in the ag qVOA were incubated for up to 72 hours to maximize T cell activation and HIV-1 expression. Following a 21-day co-culture with the MOLT-4 target cell line, exponential viral outgrowth was quantified on day 21 by p24 ELISA, and viral RNA was isolated from culture supernatants for sequence analysis.

**Figure 1.**
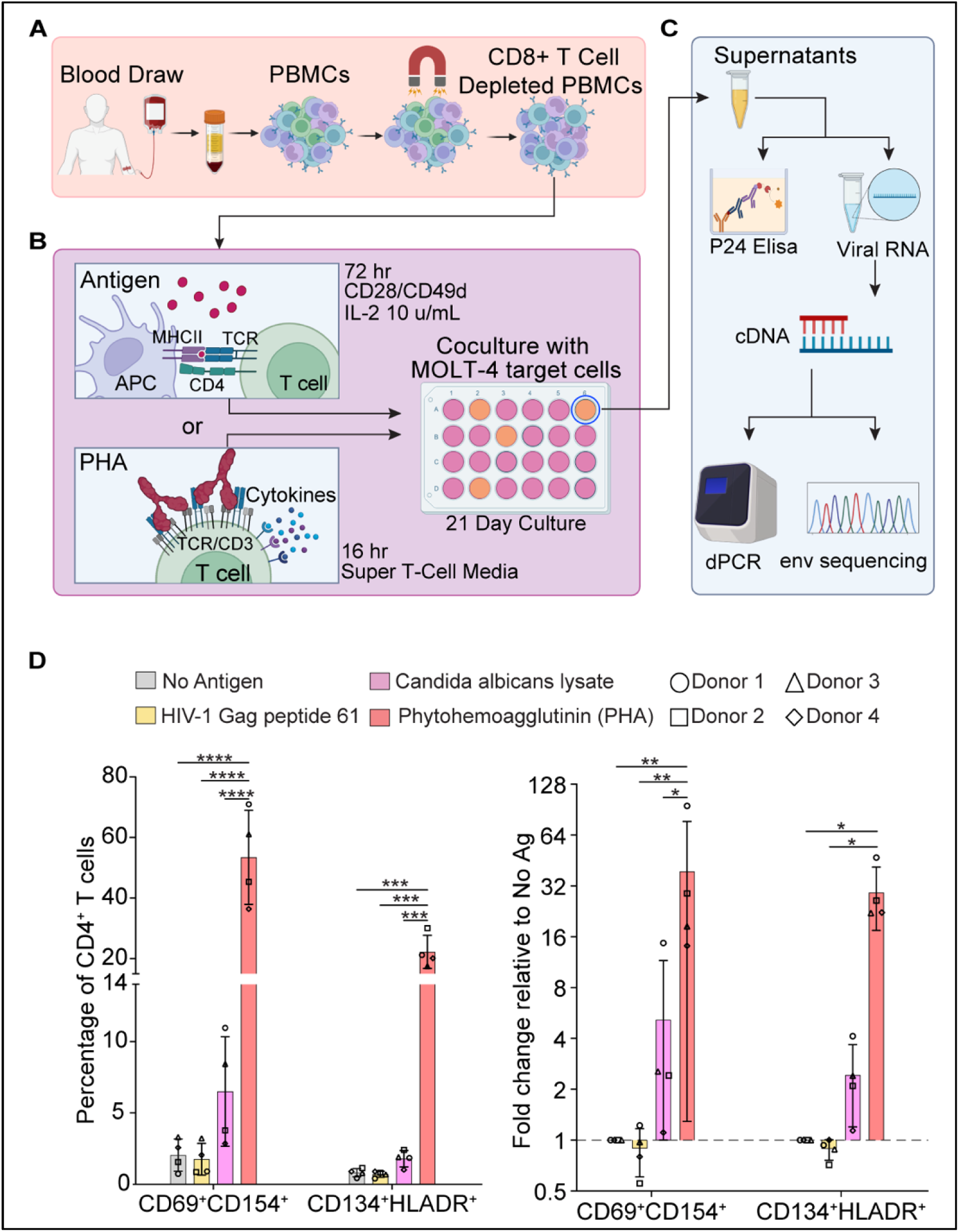
Experimental design of the Antigen Restricted Quantitative Viral Outgrowth Assay (ag qVOA). **A.** PBMCs were isolated from whole blood and depleted of CD8^+^ T cells. **B.** Cells were then stimulated with antigens or PHA and co-cultured with MOLT-4/CCR5 target cells under conditions supporting either low-level (antigen) or robust (PHA) T-cell activation. Cultures were maintained for 21 days. **C.** HIV-1 outgrowth in culture supernatants was measured by p24 ELISA; positive wells were used to calculate infectious units per million (IUPM) and for sequencing of virion-associated HIV-1 RNA. **D.** Expression of activation-induced markers at 24 hours of culture in CD8-depleted PBMCs from HIV-1 negative donors stimulated with no antigen (No Ag), HIV-1 Gag peptide 61 (P61), Candida albicans lysate (CA lysate), or phytohemagglutinin (PHA). Each data point represents the average of 2 technical replicates from an individual donor: n = 4 donors. Bars represent mean ± SD. *p < 0.05, **p < 0.01, ***p < 0.001, ****p < 0.0001 by two-way ANOVA with Tukey’s multiple comparisons test. See also Figure S1.

To validate activation specificity, we stimulated CD8-depleted PBMCs from four HIV-1-negative donors with no antigen, the HIV-1 Gag peptide P61 (STLQEQIGWMTNNPP), a protein lysate from the opportunistic pathogenic yeast *Candida albicans* ^38^ (CA), or PHA (Figure 1D). At 24 hours, PHA stimulation induced strong CD4⁺ T cell activation, characterized by high expression of activation marker pairs CD154/CD69 and CD134/HLA-DR^39,40^. The no-antigen (No-Ag) control exhibited low background activation, confirming that the culture conditions were not inherently activating. Activation by Gag P61 was comparable to the No-Ag control. While not statistically significant, CA lysate induced a modest increase in activated CD4⁺ T cells, consistent with the variable and low frequency of CA-reactive cells in peripheral blood^41^.

### Participant P012 harbors an intact provirus integrated within a pericentromeric region in an expanded *Candida albicans* responsive clone

Participant 012 (P012) is a 46-year-old African American male living with HIV-1 who initiated ART during chronic infection 18 years prior to this study. After intermittent ART adherence between years 6 and 12 following initiation, he maintained sustained adherence and viral suppression for the 6 years preceding sample collection (Figure 2A, Figure S2A). As part of a study to identify CD4^+^ T cells responsive to the ubiquitous commensal fungus CA (Camilo-Contreras, manuscript in preparation), we stimulated CD8-depleted PBMCs with CA lysate and sorted responding cells based on the upregulation of the activation-induced markers CD69^+^ and CD154^+^ (Figure 2B, Figure S2B). CA-responsive cells were enriched in total and intact HIV-1 DNA relative to non-responding cells (NR) and non-specifically stimulated cells (αCD3/CD28) (Figure 2C). This enrichment was driven by a highly expanded proviral variant comprising 47% of HIV-1 *envelope* (*env*) proviral sequences recovered from CA-responsive cells and present at 937 copies per million CA-responsive cells (Figure 2D-F).

**Figure 2.**
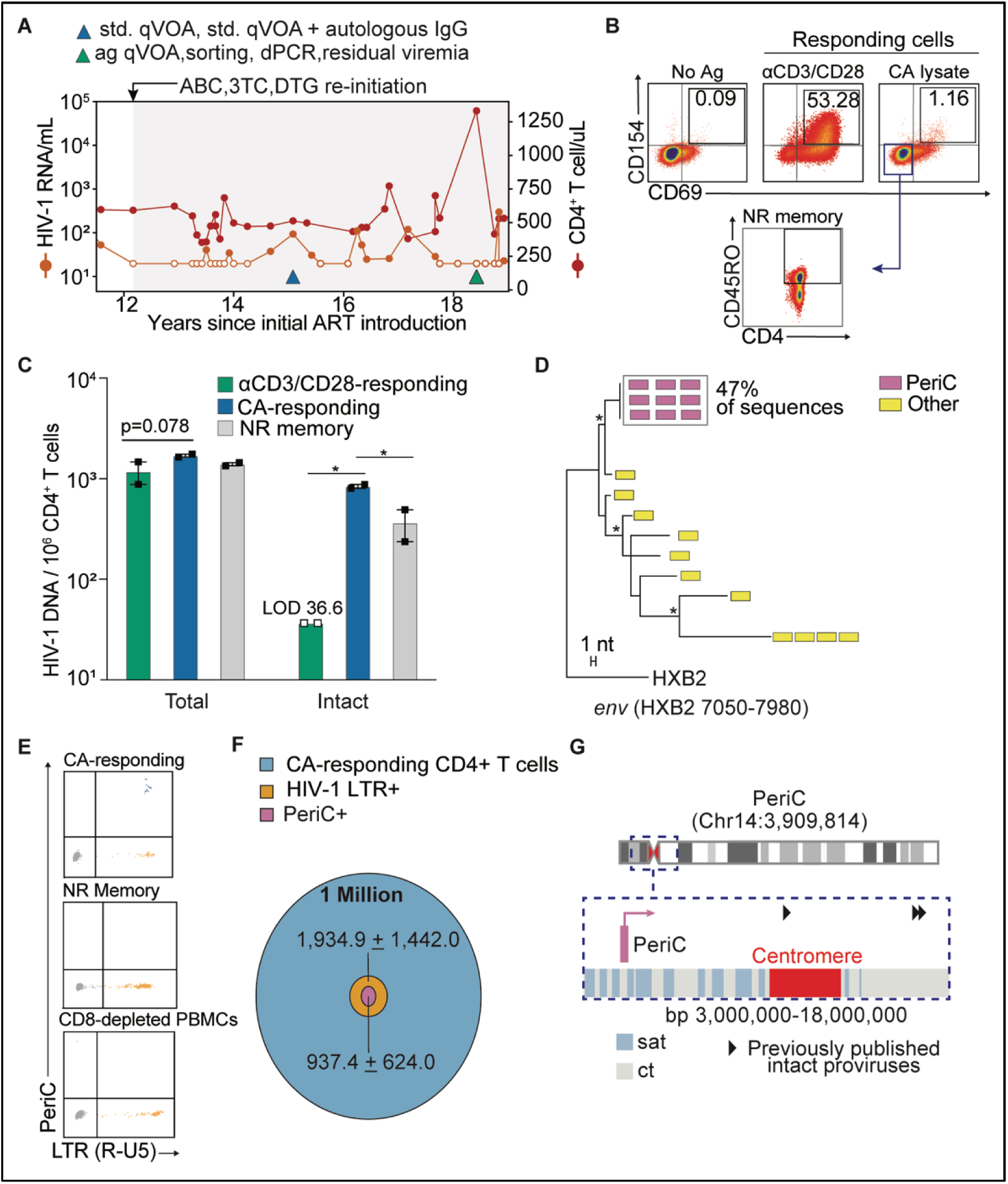
Participant P012 harbors an intact provirus integrated within a pericentromeric region in an expanded *Candida albicans* responsive clone. **A**. Longitudinal HIV-1 RNA and CD4^+^ T cell counts for P012. See also Figure S2A. **B.** Flow cytometry plots of CD8-depleted PBMCs after a 16 hr stimulation with No-Ag, αCD3/CD28 or CA lysate, gated on live CD4^+^ T cells. Gating for CA non-responding memory cells is shown. Also see Figure S2B. **C.** Total and intact HIV-1 DNA copies per 10^6^ CD4^+^ T cells in sorted cells, as measured by IPDA. Bars represent mean ± SEM. p* < 0.05 by two-way ANOVA with Tukey’s multiple comparisons test. **D.** Neighbor-joining (NJ) phylogenetic tree of HIV-1 *env* sequences from CA-responding cells. **E-F.** dPCR quantification of LTR and PeriC copy numbers in CA-responding, NR memory, and CD8-depleted PBMC populations. **G.** Integration site and relative orientation of PeriC on chromosome 14 (Chr14:3,909,814; T2T-CHM13v2.0/hs1). Insert shows the centromeric region with the PeriC integration site (pink) and previously published intact proviruses (black triangles). Centromeric satellite DNA (sat) and centromeric transition regions (ct) are indicated. See also Figure S2C. Open symbols indicate values below the limit of detection: <20 cp/mL (**A**), 36.6 copies/10□ CD4⁺ T cells (**C**).

Paired proviral and integration site analysis^42,43^ revealed that this provirus is genetically intact and located within a centromeric transition (ct) region on the p arm of chromosome 14 (Chr14: 3,909,814, based on the T2T human genome assembly): a pericentromeric domain of a mixture of satellite DNA and other sequence elements flanked by annotated satellite (sat) regions, hereafter referred to as “PeriC” (Figure□2G). Chromosome□14 is acrocentric, with a short p arm enriched in repetitive ribosomal RNA gene clusters and largely devoid of protein-coding genes^44^. RefSeq annotations confirmed the absence of nearby annotated genes, Precision Run-On Sequencing (PRO-seq) from unstimulated and PMA/ionomycin-stimulated CD4⁺ T cells showed no detectable nascent RNA expression in this region^45,46^, and epigenetic profiling using Epilogos indicated a quiescent chromatin state consistent with transcriptional inactivity^46,47^ (Figure S2C). Notably, intact proviruses in this region have been found in ECs^26^ and in individuals on ART^34,48^ (Figure 2G).

### The pericentromeric provirus PeriC can be reactivated by cognate antigen and contributes to residual viremia

P012 was previously extensively characterized by Bertagnolli et al. using a large-scale qVOA with or without the addition of autologous IgG^49^ (Figure 2A), in which PeriC represented a minor fraction of outgrowth sequences (7/97) and was sensitive to autologous neutralizing antibodies (aNAb). To test cognate antigen-driven reactivation, we performed the ag qVOA using CA lysate, PHA, or no-antigen. Exponential viral outgrowth was observed in both stimulated conditions, and none in the no-antigen control (Figure 3A). The PHA condition yielded more wells with p24 levels above the limit of detection (100 pg/mL), possibly reflecting improved MOLT-4 viability and HIV-1 infectivity in the cytokine-rich Super T Cell Media used in that condition (data not shown). CA lysate stimulation produced multiple viral variants in supernatants at 72 hours, but only PeriC drove exponential outgrowth in the ag qVOA (Figure S2E-F), suggesting the other variants represented defective RNA packaged into virions during the initial stimulation. Of note, supernatants alone did not initiate outgrowth in MOLT-4 cells (Figure S2G).

**Figure 3.**
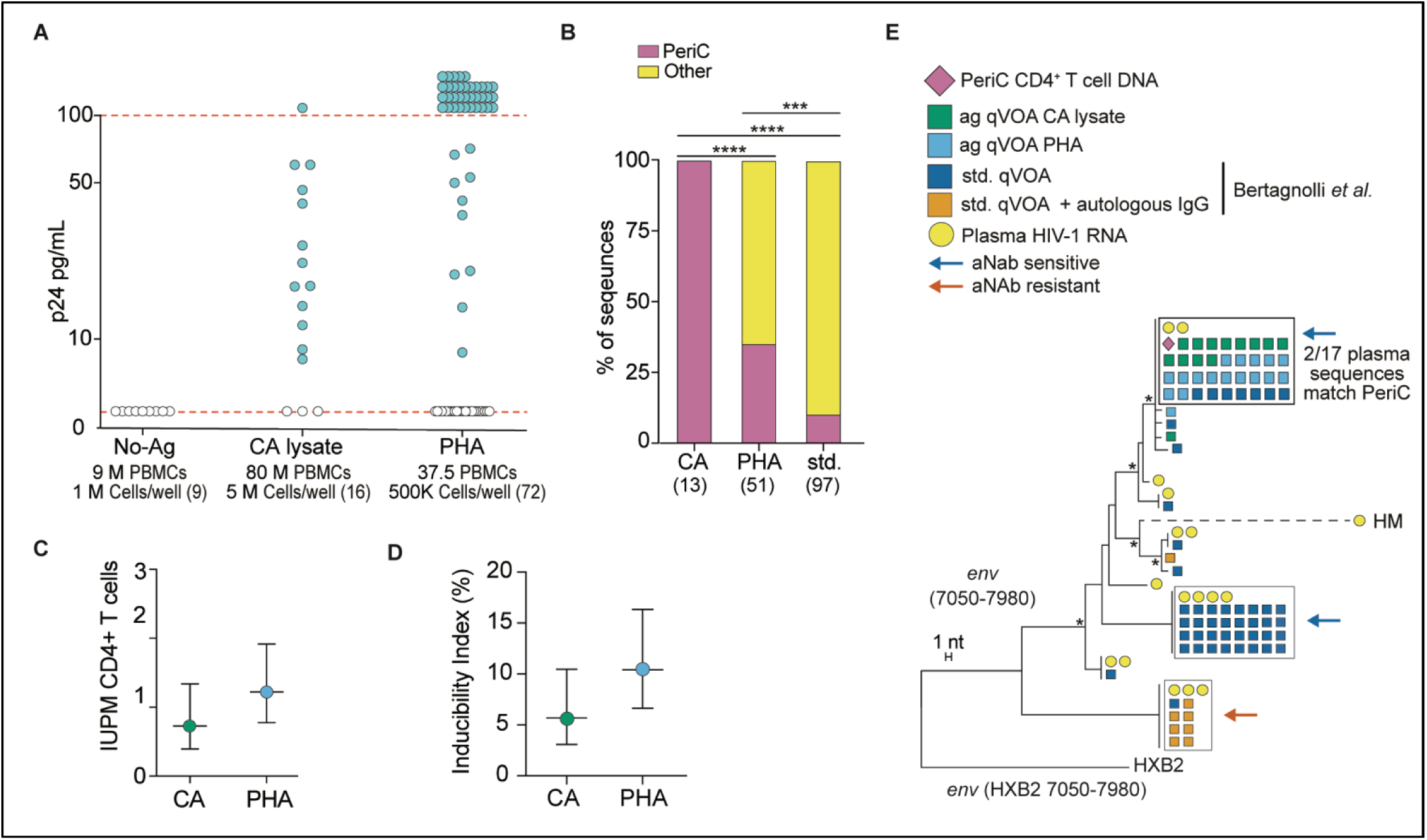
A provirus in a pericentromeric region is reactivated by cognate antigen and contributes to residual viremia. **A.** p24 pg/mL in ag qVOA culture supernatants on day 21. The total number of wells is indicated in parentheses. Open symbols indicate values below the limit of detection: 5 pg/mL. **B.** Proportion of sequences matching PeriC. Numbers in parentheses indicate number of sequences. ***p < 0.001, ****p < 0.0001 by or Fisher’s exact test **C.** PeriC-specific IUPM and **D.** inducibility index in CD4⁺ T cells following CA lysate or PHA. CD4^+^ T cell frequency was determined by flow cytometry analysis (see also Figure S3). Bars represent the maximum likelihood estimate; error bars indicate 95% confidence intervals. **E.** NJ tree of HIV-1 *env* sequences from cellular DNA, std. qVOA/ag qVOA outgrowth virus, and plasma RNA during ART. The std. qVOA was performed with or without autologous IgG in Bertagnolli *et al.*; orange and blue arrows indicate aNAb-resistant and aNAb-sensitive outgrowth virus, respectively. Only sequences closely related to plasma variants are shown; for all outgrowth sequences see also Figure S2D.

CA lysate-driven outgrowth was significantly enriched for PeriC, with 100% of recovered *env* sequences matching this provirus, compared to 35% in the PHA-stimulated condition and 10% in the std. qVOA from Bertagnolli *et al* (Figure 3B, S2D), consistent with antigen specific reactivation in the ag qVOA. To estimate reactivation efficiency, we calculated PeriC-specific infectious units per million (IUPM) and calculated an inducibility index from the number of PeriC proviral copies plated per well and the wells with detectable PeriC outgrowth. PeriC inducibility was only modestly higher following PHA stimulation (9.54%) than CA lysate stimulation (5.69%) (Figure 3C-D).

To assess whether PeriC contributed to residual viremia in vivo, we analyzed P012 plasma HIV-1 RNA. Although viremia was undetectable by clinical assay at the time of sampling (Figure 2A), 4 copies/mL were detected from 2 ml of plasma by ultrasensitive dPCR assay targeting the 5’Leader region^50^ (Figure S2H). Plasma-derived *env* sequences were largely oligoclonal, with several matching sequences recovered from the std. qVOA (Figure 3E). Importantly, two of 17 sequences matched PeriC, indicating this provirus contributes to residual viremia during suppressive ART despite its genomic location being unfavorable for HIV-1 gene expression.

### An Elite Controller with a ZNF Gene Provirus in an HIV-1 Gag-responsive Clone

We next examined a second study participant ES24, a 68-year-old African American male. ES24 is an HLA-B*57^+^ EC who has maintained undetectable viremia in the absence of ART for over a decade and has been previously characterized by our group^17^. Five years prior to this study, he initiated ART to be treated for metastatic lung cancer with chemotherapy combined with radiation (CRT), and anti-PD-L1 therapy (Figure 4A). At this time, his reservoir was dominated by two intact proviruses integrated into zinc finger (ZNF) genes, termed ZNF470i and ZNF721i. ZNF gene exons are enriched for the repressive histone modification H3K9me3^51^, and intact proviruses integrated into ZNFs are selectively enriched in ECs and individuals on LT-ART, consistent with a role for epigenetic silencing in immune evasion^26,28,34,52^. Importantly, *ZNF721* is the ZNF gene with the most intact proviruses recovered across PWH to date.

**Figure 4.**
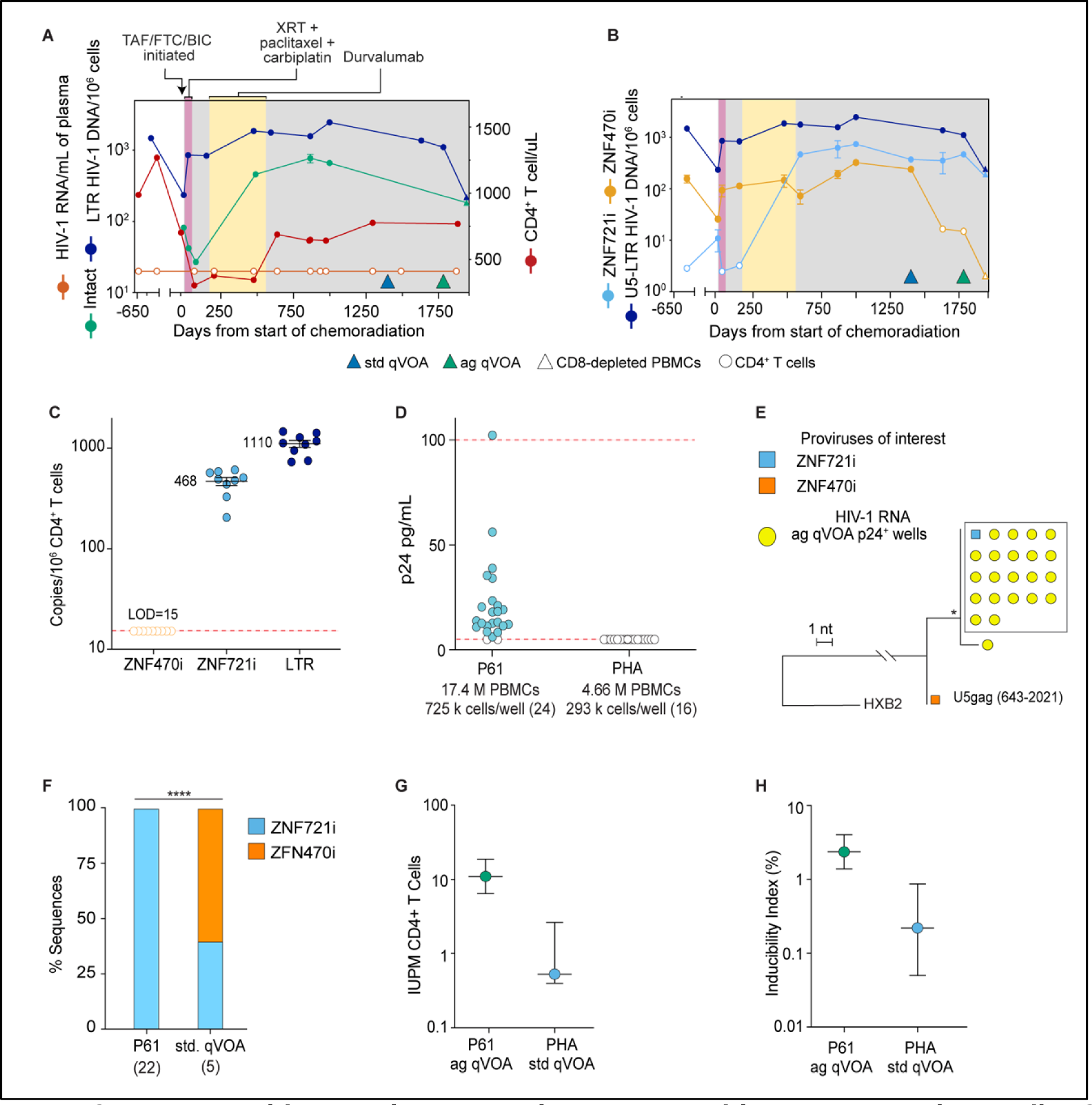
Cognate Peptide Reactivates Provirus Integrated into ZNF gene in an Elite Controller. **A.** Longitudinal plasma HIV-1 RNA and CD4^+^ T cell counts for ES24. Open circles denote undetectable plasma viremia (<20 copies/mL). Quantifications of HIV-1 LTR and intact HIV-1 DNA are also shown. Error bars represent SEM values. **B.** Longitudinal dPCR quantification of LTR and proviruses of interest. Open symbols indicate values below the limit of detection. Error bars represent SEM. **C.** dPCR quantification in CD4^+^ T cells of ZNF470i, ZNF721i and LTR at the ag qVOA timepoint. Each symbol represents a dPCR reaction, and a total of 274K CD4^+^ T cells were screened. Open symbols represent measurements below the limit of detection. Bars represent mean ±SEM. **D.** p24 ELISA values from day 21 of ag qVOA cultures. Open symbols represent measurements below the limit of detection. Number in parenthesis represents number of wells. **E.** NJ phylogenetic tree of HIV-1 U5-gag sequences. **F.** Proportion of outgrowth sequences from ag qVOA and std. qVOA matching the proviruses of interest. No sequences were recovered from the PHA ag qVOA condition. Number in parenthesis represents number of sequences. *p < 0.05, ***p < 0.001, ****p < 0.0001 by Fisher’s exact test. **G**. ZF721i specific IUPM (left) and inducibility index (right) in P61 ag qVOA and std. qVOA. Bars represent the maximum likelihood estimate; error bars indicate 95% confidence intervals. CD4^+^ T cell frequency was determined by flow cytometry analysis (see also Figure S3).

ZNF470i initially represented the dominant intact proviral clone and tracked with the overall contraction and expansion of infected cells during treatment but became undetectable by day 1632 post-CRT. ZNF721i, initially a minor clone, progressively expanded to become the dominant intact provirus (Figure 4B-C). In our previous study^17^, both ZNF470i and ZNF721i exhibited low inducibility in the std. qVOA (0.1-1%), consistent with transcriptional repression at their integration sites. Notably, ZNF721i resides within a clone responsive to an HIV-1 Gag peptide (peptide 61), enabling antigen-specific ex vivo stimulation at the peptide level. Upon cognate peptide stimulation, ZNF721i produced approximately 200-fold less virus than a separate Gag-reactive defective provirus (Chr7.d11sc), and ZNF721i-carrying cells proliferated ex vivo with limited CD8⁺ T cell-mediated clearance^17^.

### A cognate HIV-1 Gag peptide induces viral outgrowth despite integration into ZNF721

To assess the antigen-specific reactivation of ZNF721i, we performed the ag qVOA on cells collected on day 1785 post-CRT using Gag peptide 61 or PHA as a non-specific control. No outgrowth was detected with PHA stimulation (Figure 4D), which was consistent with the low ZNF721i inducibility in the std. qVOA (0.22%) and the limited number of ZNF721i proviruses plated in that condition (675 copies). Peptide 61 stimulation resulted in exponential viral outgrowth, and all 22 U5*gag* RNA sequences recovered from p24+ wells matched ZNF721i, confirming that cognate antigen engagement was sufficient to reactivate this provirus despite its transcriptionally repressive integration site (Figure 4E).

Given the absence of outgrowth in the ag qVOA PHA condition, we used the previously conducted std. qVOA to compare antigen-specific and polyclonal reactivation. ZNF721i was significantly enriched in the peptide 61 ag qVOA (22/22 sequences) relative to the std. qVOA (2/5 sequences), consistent with antigen-specific reactivation (Figure 4F). Peptide 61 stimulation also yielded a higher IUPM and an approximately tenfold higher inducibility index (2.42% vs 0.22 %) compared to the std. qVOA (Figure 4G-H).

## DISCUSSION

Intact proviruses integrated within genomic regions less favorable for HIV-1 transcription become progressively enriched during long-term ART and in ECs, yet the degree of such “deeper latency” and their contribution to viral rebound remain incompletely defined. Here, we demonstrate that intact proviruses integrated within two genomic contexts associated with transcriptional repression—a pericentromeric region and a ZNF gene—can be selectively reactivated through cognate antigen stimulation, providing insight into mechanisms that may drive viral reactivation in vivo.

To directly test antigen-driven reactivation, we applied the ag qVOA to two deeply characterized individuals^17,49^. In both participants, proviruses of interest were selectively reactivated by their cognate antigen and highly enriched among outgrowth sequences relative to polyclonal stimulation conditions. These findings confirm that the ag qVOA achieves antigen-specific latency reversal without significant non-specific activation. Beyond this study, the ag qVOA provides a framework to evaluate latency reversal within antigen-defined T cell populations and could be extended to assess whether common co-infections (e.g., *mycobacterium tuberculosis,* cytomegalovirus) contribute to reservoir enrichment within pathogen-reactive CD4^+^ T cell populations^53–55^.

PeriC and ZNF721i exhibited distinct inducibility profiles. PeriC showed measurable inducibility following both antigen-specific and polyclonal stimulation and contributed to residual viremia in vivo. This may reflect a pericentromeric chromatin environment that, despite being heterochromatic, permits proviral transcription, as recently seen in other pericentromeric proviruses^34^. In contrast, ZNF721i exhibited minimal inducibility with PHA but an approximately tenfold increase following cognate antigen stimulation, suggesting that TCR-peptide-MHCII signaling can overcome transcriptional constraints at this locus. The different inducibility between the pericentromeric and the ZNF proviruses described here is in agreement with recent work from Bittar^34^ and Ferreira^56^ on reservoir clones isolated from PWH by limiting dilution and expansion in vitro. Collectively, these studies show higher HIV-1 transcription and p24 expression from proviruses in centromeric or pericentromeric regions than in ZNF genes. The mechanisms underlying these differences remain unclear but may involve differences in transcription factor accessibility and RNA Pol II-dependent transcription between these two chromatin environments, contributing to heterogeneous inducibility among proviruses classified within repressive chromatin.

The in vivo expression of PeriC and ZNF721i further reflect differences in chromatin environment, antigen exposure, and immune pressure. PeriC-derived sequences were detected in residual viremia despite integration within repressive chromatin, suggesting that frequent commensal antigen encounter sustains in vivo expression. Of note, PeriC contributed to viremia despite aNAb sensitivity^49^, suggesting that aNAbs can block viral outgrowth but cannot completely mediate immune clearance of virus-expressing cells^57^. In contrast, no HIV-1 RNA was detected in ES24’s plasma when screening 2 mL by dPCR or 5 mL using single genome U5*gag* sequencing. The lack of ZNF721i detection in plasma despite its higher frequency in peripheral CD4^+^ T cells is consistent with the lower ZNF721i inducibility, limited antigen availability, and potent CD8^+^ T cell-mediated killing of HIV-1-expressing cells, previously demonstrated in this participant^17^. Notably, Halvas *et al.* showed that integration in ZNF genes do not confer absolute transcriptional silencing, as intact proviruses in *ZNF721* and *ZNF268* contributed to non-suppressible viremia in two PWH and could be induced by std. qVOA^52^.

Collectively, these findings refine current models of HIV-1 latency by demonstrating that epigenetically repressed proviruses are not uniformly inert and can remain inducible upon antigen recognition. While repressive chromatin reduces inducibility, it does not confer permanent silencing. Thus, these proviruses remain clinically relevant, contributing to residual viremia and viral rebound if resistant to immune pressure. The selection of intact proviruses in heterochromatin may serve as a marker of effective interventions aimed at enhancing the killing of reservoir cells expressing residual HIV-1^30^. However, reservoirs enriched for such proviruses may favor the success of immune-based HIV-1 cure strategies, as reduced virus expression during analytical treatment interruption may provide a window of opportunity for host cellular and humoral immunity to expand and control infection, as recently observed in post-intervention controllers^58,59^.

## LIMITATIONS

This study has several limitations. First, the analysis is restricted to two individuals, reflecting the rarity of identifying intact proviruses within clonally expanded CD4⁺ T cells of known antigen specificity. While this constrains generalizability, the depth of characterization provides insight into antigen-driven reactivation from repressive chromatin contexts. Second, longitudinal proviral quantification was not available for the individual on long-term ART, precluding direct assessment of temporal changes in proviral dynamics. Importantly the ag qVOA is an ex vivo system involving high antigen concentrations and may not fully recapitulate the complexity of antigen exposure, tissue localization, and immune surveillance in vivo. Finally, the contribution of additional antigen specificities and chromatin environments to antigen-driven reactivation remains to be determined in larger cohorts. Future studies extending this approach to additional individuals and antigens will be necessary to establish the broader relevance of these findings.

## METHODS

### Sex as a biological variable

Due to the rarity of the specific proviruses investigated in this study, the study was limited to two male individuals. Consequently, sex was not evaluated as an independent biological variable, and the study was not designed or powered to assess sex-based differences in antigen-driven HIV-1 proviral reactivation. While the mechanisms examined are not expected to be exclusive to males, the generalizability of these findings to females cannot be determined from the current study. Future studies including larger cohorts with both male and female participants will be necessary to evaluate the influence of sex and to establish the broader applicability of these findings.

### Study participant ES24

ES24 is a 68-year-old African American male who was diagnosed with HIV-1 15 years prior to this study and has maintained consistent viral suppression (<20 copies/mL) since. Whole blood was collected from the participant, and PBMCs were isolated using Ficoll-based density gradient centrifugation. CD8⁺ T cells were depleted using magnetic bead-based separation (Miltenyi Biotec).

### Study participant P012

P012 is a 45-year-old African American male who was diagnosed with HIV-1 24 years prior to the study. Coded, de-identified leukapheresis products were shipped by same-day courier to the Johns Hopkins School of Medicine for processing and analysis under a protocol approved by a Johns Hopkins University School of Medicine IRB. PBMCs were obtained from cryopreserved leukapheresis samples, rested for 2 hours in RPMI (Gibco) media containing 10% heat inactivated fetal bovine serum (FBS).

### MOLT-4/CCR5 cell line and cell culture

MOLT-4/CCR5 is a human T-lymphoblastoid cell line engineered to stably express CCR5. The cell line was obtained from the NIH AIDS Reagent program (ARP-4984) and was cultured in suspension in RPMI□+□10 % FBS□+□1% PenStrep.

### Antigens

CA Lysate: *C. albicans* lysate was prepared from strain ATCC90028. The fungus was cultured in yeast extract peptone dextrose media (YPD) at 30□°C to stationary phase, then expanded in YPD supplemented with 10% heat-inactivated FBS at 37□°C for 5 hours. Cells were collected, washed in sterile PBS, and frozen at -80°C. After thawing, cells were lysed using a Glen Mills G-M French Press (5 passes at 25,000 PSI), clarified by centrifugation, and sterile-filtered (0.22□µm polyvinylidene fluoride). Protein concentration was determined by Bicinchoninic acid (BCA) assay, and lysates were stored at –80□°C until use at 20□µg/mL. An optimal stimulation concentration of 20□µg/mL was determined by titration using cells from uninfected donors. Gag P61: Gag peptide 61 (STLQEQIGWMTNNPP, corresponding to Gag 241-255) was obtained from NIH HIV Reagent Program and used at 10□µg/mL.

### Sorting of CA-responding, non-responding memory, and anti-CD3/CD28-responding CD4⁺ T cells

CD8-depleted PBMCs from P012 were cultured in RPMI (Gibco) supplemented with 10% human serum, anti-CD40 (1□µg/mL, Miltenyi), CD28/CD49d (0.5□µg/mL, BD Biosciences), and antiretroviral drugs (DTG, FTC, TDF). Cells were stimulated with either no treatment, anti-CD3/CD28 beads, or CA lysate (20□µg/mL). After 16 hours, responding CD4⁺ T cells were identified by upregulation of activation induced markers (CD137⁺, CD154⁺, CD69⁺) and sorted using fluorescence-activated cell sorting (FACS). Non-responding memory CD4⁺ T cells were sorted from the CA lysate condition based on CD45RO expression and lack of activation marker upregulation (CD154⁻, CD69⁻, CD45RO^high^). For HIV-1 DNA characterization experiments, CD4⁺ T cell populations were sorted using a MoFlo XDP High-Speed Cell Sorter (Beckman Coulter). For PeriC provirus quantification, cell sorting was performed using the MACSQuant Tyto Cell Sorter (Miltenyi Biotec). Sorting gating strategies and antibody panels are detailed in Figure S2B and Supplementary Table 2.

### Intact proviral DNA Assay (IPDA)

Cellular guide DNA (gDNA) was isolated from CD8-depleted PBMCs, whole PBMCs, or CD4⁺ T cells as previously described^17^. IPDA was performed according to established protocols^36^ using either the QX200 Droplet Digital PCR System (Bio-Rad) or the QIAcuity One Digital PCR System (Qiagen). Longitudinal IPDA data for ES24 samples collected between days - 635–1400 post-CRT were reported in a previous publication from our group^17^ .

### Provirus sequencing

Proviral sequencing for ES24 was performed as previously described^17^. For P012, the *env* region was sequenced from genomic DNA (gDNA) by a nested PCR using Platinum Taq High Fidelity DNA Polymerase (Thermo Fisher Scientific). Outer PCR thermocycling conditions: 94°C for 2 minutes, 45 cycles of 94°C for 30 seconds, 50°C for 30 seconds, 72°C for 2 minutes, followed by 72°C for 3 minutes and a hold at 4°C (Env-outer-forward: 5’-GCCAGTAGTRTCAACYCAA-3’; Env-outer-reverse: 5’-GCARATGAGTTTTCYAGAGCA-3’). Inner PCR conditions: 94°C for 2 minutes, 45 cycles of 94°C for 30 seconds, 55°C for 30 seconds, 72°C for 2 minutes, and then 72°C for 3 minutes and a hold at 4°C (Env-inner-forward: 5’-CTGCTAAATGGCAGTCTAGC-3’; Env-inner-reverse: 5’-TTGCCTGGAGCTGYTTRATGC-3’). PCR product from positive reactions was sent for purification and Sanger sequencing (Azenta). A full-length proviral sequence for PeriC clone was obtained using matched integration site and proviral sequencing (MIPseq), with flanking LTR regions recovered using primers complementary to the human genome at the integration site and internal to the provirus as described^17^.

### Integration site analysis

Integration sites for ES24 were analyzed as previously described^17^. For P012, gDNA from CA*-*responding CD4⁺ T cells was diluted and amplified using the REPLI-g Advanced Single Cell Kit (QIAGEN) so that each reaction would contain only one or no proviruses. The integration site of the PeriC clone was then characterized using the Lenti-X Integration Site Analysis Kit (Clontech). Resulting PCR products were sequenced by Sanger (Azenta). The resulting sequences were aligned to the human genome assembly T2T-CHM13v2.0 using the UCSD human BLAT search.

### Quantification of proviruses of interest and total LTR copies by dPCR

For ES24, U5-LTR copies and specific proviruses (ZNF470i and ZNF721i) were quantified by digital PCR as previously described^17^ using either CD4⁺ T cells or CD8⁺ T cell-depleted PBMCs. Where possible, CD8-depleted PBMCs were analyzed by spectral flow cytometry (Cytek Aurora), and results were normalized to CD4⁺ T cell equivalents (gating strategy detailed in Figure S3). For P012, PeriC was quantified through digital PCR (QIAcuity One System, Qiagen) using a fluorescent probe specific to the host-U3 junction at the HIV-1 integration site^15^ (PeriC forward primer: 5’-AGAGATCCCTCAGACCCTTTAG- 3’; PeriC reverse primer 5’-GCATCTAGCAAACCAGC AATTT-3’; PeriC probe 5’- /56FAM/TGGAAAATC/ZEN/TCTAGCAAAAGGTGT TATTGG/3IABkFQ/ -3’).

### Antigen-restricted Quantitative Viral Outgrowth Assay (ag qVOA)

CD8-depleted PBMCs were stimulated with antigens for 72 hours and subsequently co-cultured with the permissive MOLT-4/CCR5 cell line (Baba et al., 2004) in minimally activating media supplemented with anti-CD40, CD28/CD49d, and IL-2 (10 U/mL). In parallel, cells were also stimulated with PHA for 24 hours and co-cultured with MOLT-4/CCR5 cells in a cytokine-rich medium containing 2% T cell growth factor and 10 U/mL IL-2, as previously described^35^. Cultures were split and the medium replenished every 72-96 hours. On days 14 and 21, HIV-1 replication in the culture supernatant was assessed using an HIV-1 p24 antigen ELISA (NEK050001KT, Revvity), and supernatants from positive wells were collected. Infectious units per million (IUPM) were calculated using IUPMStats v1.0, which implements a maximum-likelihood model as described by Rosenbloom et al. (2015). The inducibility index for each specific provirus was calculated by dividing the IUPM by the number of proviruses seeded. Although PBMCs were used as cell inputs to permit antigen presentation, IUPM and inducibility index values were normalized to CD4⁺ T cells using the CD4⁺ T cell frequency measured by flow cytometry prior to plating in the MOLT-4/CCR5 co-culture (Figure S3).

### RNA quantification and Sequencing from ag qVOA D21 outgrowth virus

Virion-associated HIV-1 RNA from positive wells was isolated as previously described^50^. HIV-1 U5-gag region was sequenced as described^17^. Similarly, the full *env* region were sequenced as described in Bertagnolli et al^49^.

### RNA quantification and Sequencing from ag qVOA stimulation culture supernatants

During a repeat experiment, stimulation culture supernatants from the CA lysate and no-antigen conditions were collected at 72 hours. Virion-associated HIV-1 RNA was isolated as described in Box *et al*^50^. To quantify polyadenylated HIV-1 RNA transcripts, complementary DNA (cDNA) was synthesized using an oligo(dT) primer and Induro Reverse Transcriptase (New England Biolabs). cDNA was subsequently quantified by digital PCR (dPCR) on the QIAcuity One Digital PCR System (Qiagen) using previously published primers and probes^60^. Due to sample availability, three culture supernatant replicates of 110 µL and one replicate of 32.5 µL were screened for HIV-1 Poly(A) for the CA lysate and no-antigen conditions, respectively. The isolated RNA was also subject to HIV-1 env cDNA synthesis and sequenced for HIV-1 *env* as described in the “Provirus sequencing” section.

### HIV-1 negative donor stimulation quantification

CD8- depleted PBMCs from uninfected donors were stimulated as described in the ag qVOA section. Cells were harvested at 24 post-stimulation for analysis by spectral flow cytometry (Cytek Aurora). The gating strategy antibody panel are provided in Figure S1 and Supplementary Table 1.

### RNA quantification and Sequencing from Plasma

Virions were isolated from plasma and RNA was extracted and quantified using the CLAWS assay as described in Box *et al*^50^. Extracted RNA was also sequenced for either the *env* or U5-gag region, as described above.

### Statistics

T test, ANOVAs, multiple comparisons, and Fisher’s exact tests were performed in GraphPad Prism (v10.2) (https://www.graphpad.com/features). p<0.05 were considered significant. Details of all statistical tests used and sample size information can be found in the corresponding figure legends.

### Study approval

Samples for this study were collected under protocols approved by either the University of Pennsylvania Institutional Review Board (IRB) (for P012) or The Johns Hopkins IRB (for ES24). Participants also provided written informed consent before enrollment.

### Data availability

Sequencing data generated from this study can be found in GenBank under accession numbers: PZ518423-PZ518533, PZ518534, and PZ518535-PZ518556. Data utilized in this study from previous publications can be found under accession numbers: MW077921-MW077975 and OR496826-OR496827.

## AUTHOR CONTRIBUTIONS

Conceptualization by A.C.C and F.RS; formal analysis by A.C.C, F.R.S and F.D.; investigation by A.C.C., F.D., H.Z., J.Z. and D.S.; project administration by F.R.S, J.N.B. and S.R.; writing- original draft A.C.C; review and editing A.C.C., F.R.S., S.R. resource and sample acquisition by A.C., D.S., J.L., J.D.S., R.F.S., L.J.M., P.T, and J.N.B.

## ACKNOWLEDGEMENTS

We thank the study participants who volunteered to take part in this study. We thank John Skinner and Nicole Barat for managing laboratory activities and logistics, and Eileen Scully, Michael Betts, and Brian Peters for support and guidance leading to this work. We also thank Sean Patro for his expertise on integration site analysis.

## SUPPLEMENTAL INFORMATION INDEX

Figures S1-S3, Tables S1-S2, and a supplemental reference.

