## Supplemental Figures and Tables for "Antigen Stimulation Reactivates HIV-1 Proviruses Despite Integration in Repressive Chromatin"

### Figure S1

### Gating strategy for activation quantification in the agQVOA

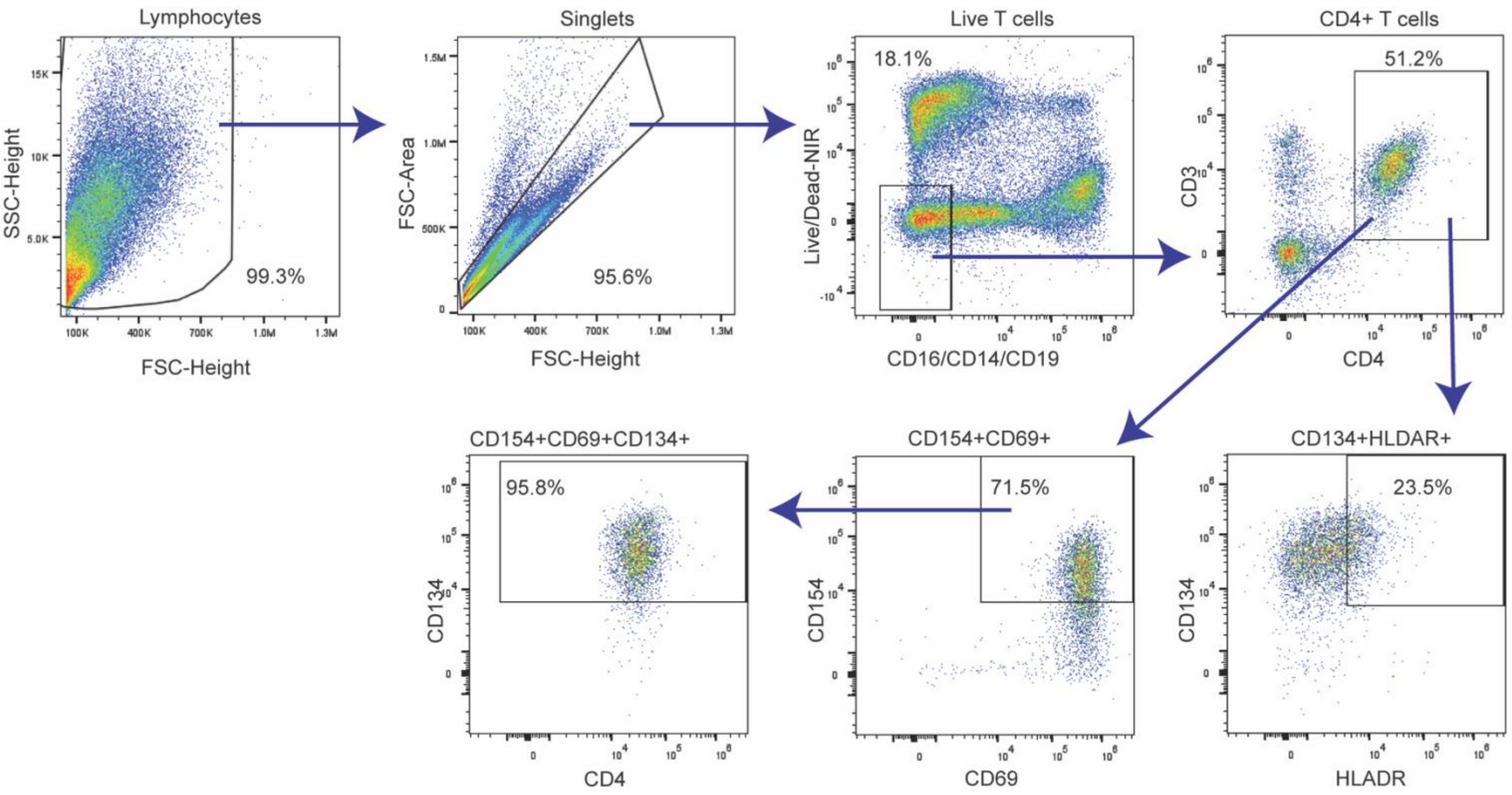

**Figure S1. Related to Figure 1. Gating strategy for activation quantification in the agQVOA**

Gating for flow cytometry analysis of activation induced marker expression in HIV negative donor PBMCs following ag qVOA stimulation. Antibody panel can be found in Table S1. Analysis was performed on a Cytek Aurora 4 Laser (16UV-16V-14B-8R).

### Figure S2

### Characterization of HIV-1 persistence and antigen-driven reactivation in participant P012

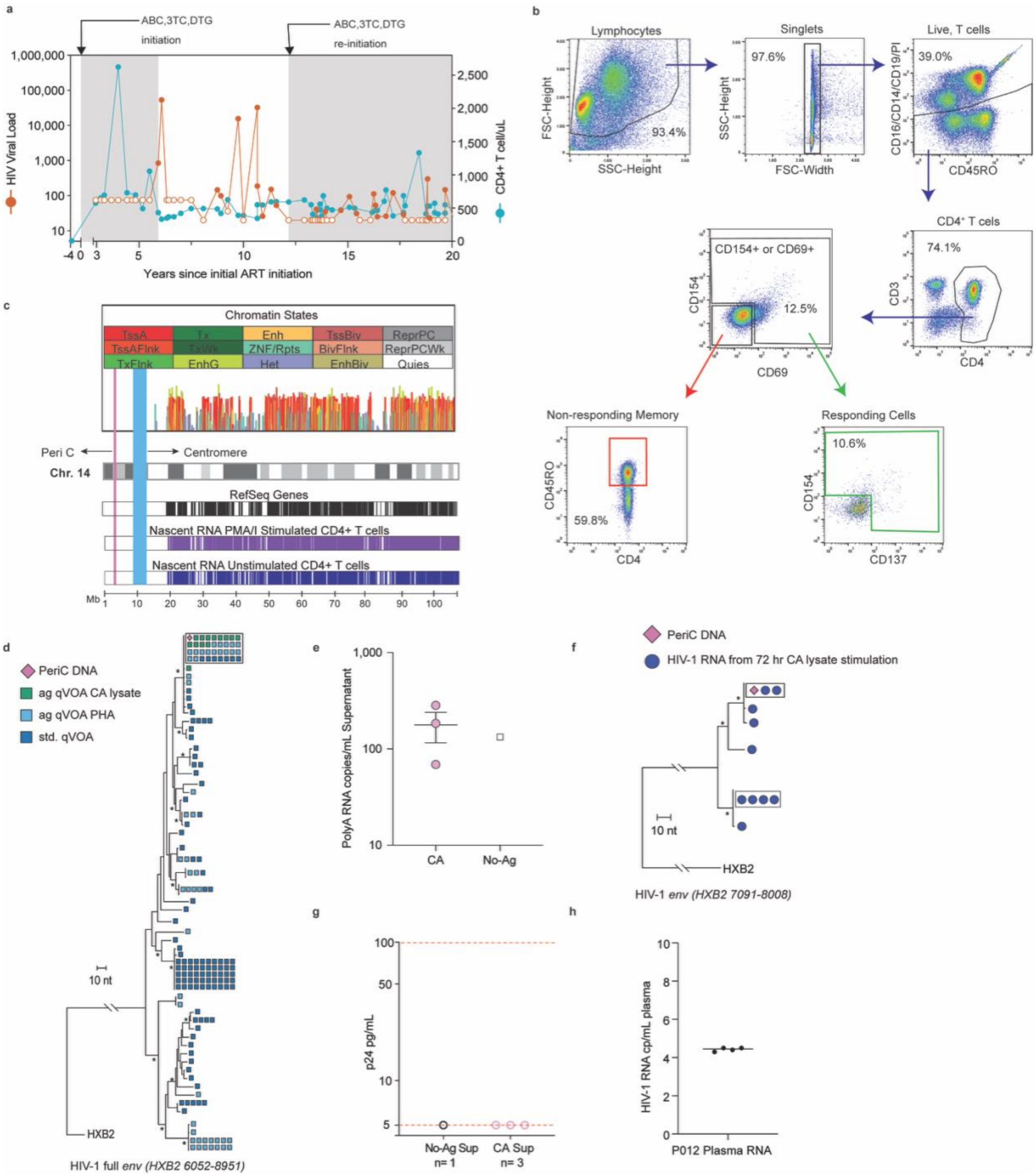

**Figure S2. Related to Figure 2 an Figure 3. Characterization of HIV-1 persistence and antigen-driven reactivation in participant P012.**

**a)** Longitudinal HIV-1 plasma viral load (left y-axis) and CD4+ T cell counts (right y-axis) for participant P012 for years -4-20 post ART initiation. Open circles represent clinically undetectable viral loads (<20 cp/mL).

**a)** Gating strategy for isolating CA-responsive, CD3/28-responsive, and non-responding memory cells in P012. Antibody panel is provided in Table S2. Cells were analyzed and sorted with a MoFlo XDP High-Speed Cell Sorter (Beckman Coulter).

**c)** Genomic features of PeriC integration site (pink vertical line) in chromosome 14. **Top:** Epigenomic chromatin state annotations across chromosome 14 in CD4+ helper T cells visualized using Epilogos (Roadmap Epigenomics, 18-state ChromHMM model, KL divergence scoring; GRCh38/hg38). Bar height reflects relative entropy of chromatin state enrichment; colors denote chromatin states as indicated in the legend. **Bottom:** Gene density and nascent transcription across chromosome 14 (GRCh37/hg19). RefSeq gene annotations are shown in black. Nascent RNA transcription (PRO-seq, minus strand) from PMA/ionomycin-stimulated (purple) and unstimulated (dark blue) primary human CD4+ T cells is shown [S1] (GEO: GSE85337).The light blue shaded region denotes the centromere.

**d)** Neighbor-joining (NJ) phylogenetic tree of HIV-1 full-length env sequences from Peri C cellular DNA, standard qVOA, and agQVOA viral outgrowth. HXB2 shown as outgroup.

**e)** HIV-1 poly(A) RNA copies per mL detected in supernatants from CA lysate-stimulated (CA) or no-antigen control (No-Ag) cultures at 72 hours. Bars represent mean ± SEM. The open square indicates values below the LOD for the No-Ag condition (<133 cp/mL).

**f)** NJ tree of HIV-1 full-length env sequences derived from PeriC cellular DNA and viral RNA in supernatant following 72-hour CA lysate stimulation in the agQVOA. HXB2 shown as outgroup.

**g)** p24 concentrations (pg/mL) in day-21 QVOA supernatants from wells containing CXCR-4 target cells only, exposed to supernatant from 72-hour CA lysate-stimulated (CA Sup, n=3) or no-antigen control (No-Ag Sup, n=1) cultures.

**h)** ddPCR quantification of HIV-1 RNA copies per mL of participants plasma at the time of agQVOA sampling. Horizontal line represents the mean.

**Figure S3**

**Flow cytometry gating strategy used to calculate CD4+ T cell percentage in ag qVOA stimulated cells.**

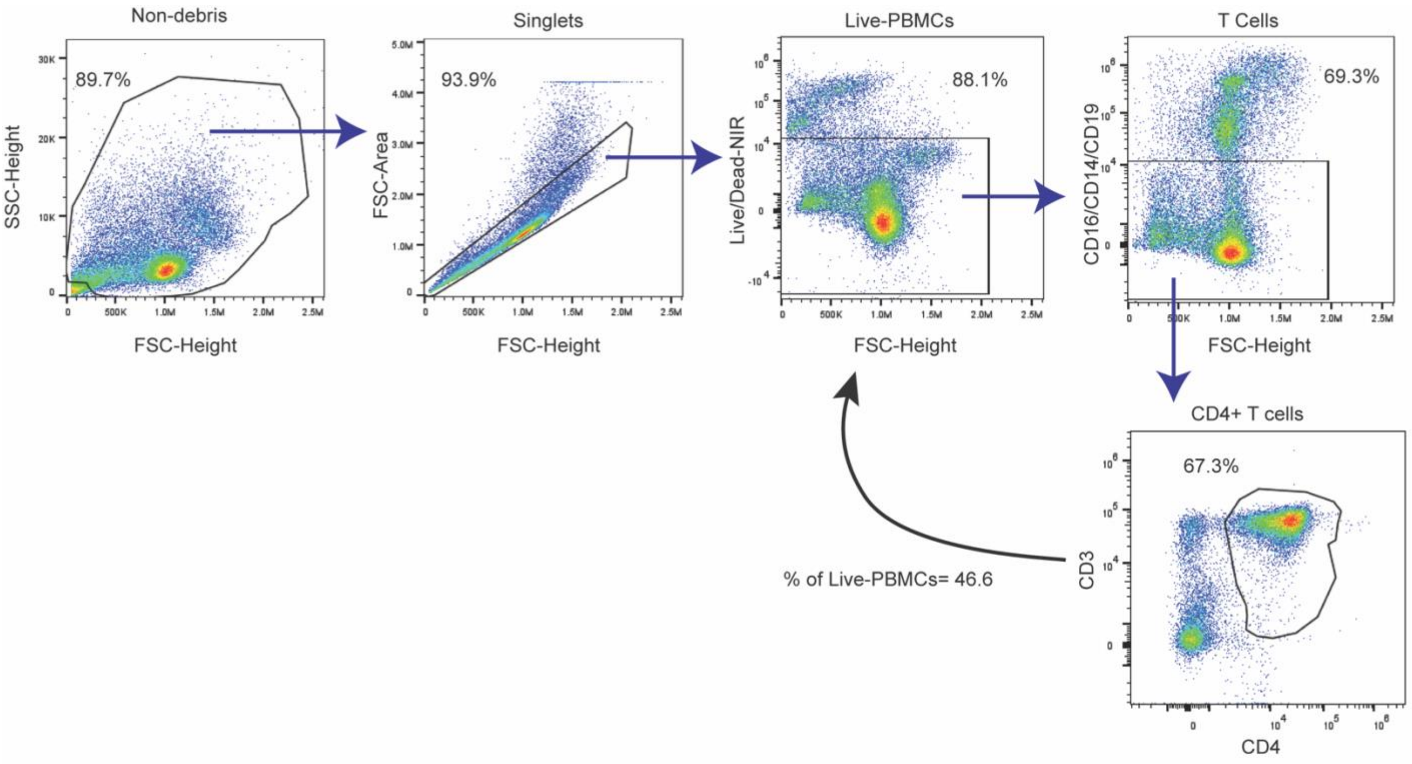

**Figure S3. Related to Forgues 3 and 4. Flow cytometry gating strategy used to calculate CD4+ T cell percentage in ag qVOA stimulated cells.**

Gating for flow cytometry analysis of percent CD4+ T cells following ag qVOA stimulation for P012 and ES24. Antibody panel can be found in Supplemental Table 2. Analysis was done on a Cytek Aurora 4 Laser 16UV-16V-14B-8R.

**Table S1**

**Antibody and viability dye panel for activation quantification in the agQVOA**

| Maker | Dye | Vendor | Clone | Catalog |
|---|---|---|---|---|
| CD4 | BUV805 | BD | OKT4 | 750976 |
| CD3 | Pacific Blue | Biolegend | OKT3 | 317313 |
| CD14 | PE-Cy5 | BioLegend | M5E2 | 301863 |
| CD16 | PE-Cy5 | BioLegend | 2H7 | 302307 |
| CD19 | PE-Cy5 | BioLegend | HIB19 | 302209 |
| CD69 | BV605 | BD | FN50 | 562989 |
| CD154 | PE/Fire™ 810 | Biolegend | 24-31 | 310857 |
| CD134 | BUV615 | BD | L106 | 751320 |
| HLADR | FITC | Biolegend | L243 | 307604 |
| Live/Dead | NIR-780 | Thermo Fisher | Viability stain | L23105 |

**Table S1. Related to Figure 1. Antibody and viability dye panel for activation quantification in the agQVOA**

Antibody and viability dye panel for cytometry analysis of activation induced marker expression in healthy donor PBMCs following agQVOA stimulation. Gating strategy can be found in Figure S1. Analysis was performed on a Cytek Aurora 4 Laser (16UV-16V-14B-8R).

**Table S2**

**Antibody panel for sorting P012 CA-responsive CD4+ T cells and analyzing CD4+ T cell percentage in ag qVOA.**

| Maker | Dye | Vendor | Clone | Catalog |
|---|---|---|---|---|
| CD4 | PE-CY7 | BioLegend | RPA-T4 | 300511 |
| CD3 | APC | BioLegend | UCHT1 | 300411 |
| CD14 | PE-CY5 | BioLegend | M5E2 | 301863 |
| CD16 | PE-CY5 | BioLegend | 2H7 | 302307 |
| CD19 | PE-CY5 | BioLegend | HIB19 | 302209 |
| CD45R0 | BV605 | BioLegend | UCHL1 | 304237 |
| CD69 | FITC | BioLegend | FN50 | 310904 |
| CD154 | BV421 | BioLegend | 24-31 | 310823 |
| CD137 | PE | BioLegend | 4B4-1 | 309803 |

**Supplementary Table 2. Related to Figure 2, Figure 3 and 4. Antibody panel for sorting P012 CA-responsive CD4+ T cells and analyzing CD4+ T cell percentage in ag qVOA.**

Gating strategy for isolating CA-responsive, CD3/28-responsive, and non-responding memory cells in P012. Gating strategy is provided in Figure S2A and S3. Cells were analyzed and/or sorted with a MoFlo XDP High-Speed Cell Sorter (Beckman Coulter) (Figures 2B, SAB) and Cytek Aurora 4 Laser (16UV-16V-14B-8R) (Figure S4).
